# Effect of compound Tinglizi Decoction on non-classical pathway of pyroptosis in rats with chronic obstructive pulmonary disease related pulmonary hypertension

**DOI:** 10.1101/2022.02.04.479096

**Authors:** Qizhi Wang, Min Liu, Yu Liu, Zhengping Bai

## Abstract

**Objective:** To observe the effect of compound Tinglizi Decoction on non-classical pathway of cell death in rats with pulmonary hypertension, and to explore its mechanism of action in the treatment of COPD-related pulmonary hypertension.

**Methods:** 60 male SD rats were randomly divided into Western group, normal group, model group, high-dose group, medium dose group and low-dose group, with 10 rats per group. We detected the protein expression levels of caspase-11, GSDMD and IL-6 by Western blot, detected serum TNF-α by ELISA, and observed the morphology of lung tissue by hematoxylin and eosin (HE) staining.

**Results:** In the expression of caspase11, GSDMD, IL-6 mRNA and protein in lung tissue, the model group was significantly higher than the normal group (P < 0.01), and the high-dose group, middle-dose group, low-dose group and western medicine group were significantly lower than the model group (P < 0.01). The levels of serum TNF-α and serum creatinine in the model group were significantly higher than those in the normal group (P < 0.01); and those in the high-dose group, middle-dose group, low-dose group and western medicine group were significantly lower than those in the model group (P < 0.05).

**Conclusion:** The mechanism of compound Tinglizi Decoction in the intervention of COPD-related pulmonary hypertension rats is related to the inhibition of non-classical pathway of cell death.

## Introduction

Chronic obstructive pulmonary disease (COPD) is a chronic, progressive respiratory disease with high morbidity and is characterized by persistent airflow limitation. In recent years, with the in-depth research on molecular biology of pulmonary hypertension, pyroptosis plays an important role in respiratory diseases(1, 2). COPD-related pulmonary hypertension (PAH) is a disease or pathophysiological syndrome of abnormally elevated pulmonary arterial pressure caused by respiratory diseases and/or hypoxia. It has pulmonary circulation disorders and a high load on the right heart, which eventually leads to right heart failure and even death. The hemodynamic criteria for pulmonary hypertension are: sea level, resting, mean pulmonary artery pressure (mPAP) 25mmHg (1mmhg = 0.133kpa) (3). The gold standard of COPD related pulmonary hypertension is right heart catheterization. Currently, the treatment method is oxygen therapy for patients with primary disease and hypoxia.

Pyroptosis is dependent on the activation of Caspase-1 and is a newly discovered form of programmed cell death in recent years. Its characteristic is that the cell continues to expand until the cell membrane ruptures, causing the release of intracellular material, which then activates a strong inflammatory response (4). When pathogens invade, the body promotes the formation of inflammasome in a certain way. This in turn activates the non-canonical caspase-4/5/11-dependent pathway and/or the canonical caspase-1-dependent pathway. In the nonclassical pathway, bacterial lipopolysaccharide (LPS) can activate caspase-4 / 5 / 11 and combine with it to induce Pyroptosis. For one thing, activated caspase-4 / 5 / 11 can activate NLRP3 inflammasome to activate caspase-1 by exposing the N-terminal domain of GSDMD protein, and then induce cell death. On the other hand, activated caspase-4/5/11 can activate the pannexin-1 channel and open the purine P2X7 channel. This promotes the release of inflammatory mediators and the efflux of intracellular K+, which activates the NLRP3 inflammasome and promotes the outward release of IL-1β. LPS can also bind to the card domain of caspase-1 to further regulate NLRP3 expression and IL-1 secretion and activation (5). In the previous experimental research of the research group, it was found that compound Tingli Zi caspule can significantly relieve the clinical symptoms and the number of acute attacks in patients with COPD-PAH. And further animal experiments confirmed that it can reduce pulmonary hypertension, improved pulmonary vascular remodeling (6–8).However, the anti-inflammatory mechanism of compound Zhixiezi decoction on COPD-PAH and whether the non-classical cell death pathway mediates its anti-inflammatory mechanism are unclear. Therefore, the purpose of this study was to observe the effect of compound Tinglizi Decoction on COPD-PAH and to explore the mechanism of its intervention in COPD-PAH.

## Materials and methods

### Animals

Sixty healthy male SD rats of SPF grade, weighing (200 ± 20) g, were purchased from Hunan Shrek Jingda Laboratory Animal Co., Ltd. with the production license number of syxk (Xiang) 2019-0017. Before the experiment, the animals were reared in the animal room for 7 days at (22 ± 2) °C. The natural circadian rhythm of light, free water intake, regular replacement of bedding. The whole animal experiment scheme was approved by the ethics committee of Hunan Institute of traditional Chinese medicine. And the disposal of experimental animals during the experiment conformed to the “Guiding Opinions on Treating Laboratory Animals Kindly” issued by the Ministry of Science and Technology of the People’s Republic of China in 2006.

### Drugs

Compound Tinglizi decoction is composed of lepidium seed 15g, peach kernel 10g, carthamus tinctorious 10g, leech 6g, ligusticum wallichii 15g, poria cocos 15g, cassia twig 15g, bighead atractylodes rhizome 15g, ardisia 10g, Scutellaria baicalensis 10g, liquorice 6g and licorice 6g. All the pieces of traditional Chinese medicine were purchased from the Chinese pharmacy of Hunan Academy of traditional Chinese medicine. These medicines were identified by Tian Qixue, chief pharmacist of Hunan Academy of traditional Chinese medicine Affiliated Hospital. and meet the relevant standards of Pharmacopoeia of the people’s Republic of China. Decocting method: compound Tinglizi decoction was added 10 times of distilled water and soaked in cold water for 30 min., then heated to boiling and decocted for 40 min. Then filter the decoction, add 8 times of distilled water, decoct as before, and filter out the decoction. The filtrate was combined twice and precipitated. After cooling, the filtrate was stored at 4 °C and used up within 1 week.

### Main reagents

Agarose (biowest, Spain, art. No. 111860); PCR loading buffer 6x (cw0610, Beijing, China); MRNA reverse transcription Kit (cw0610, Beijing, China); MiRNA reverse transcription Kit (cw2141, Beijing, China); EDTA (China Dalian Meilun Article No.: mb2514); Tris (American sigma Article No.: v900483); Trizol (thermo, USA, art. No. 15596026); Ultrasybr mixture (Kangwei century, Beijing, China, Article No.: cw2601); DEPC (sigma, USA); DM2000 plus DNA marker (cw0632, Beijing, China); Nucleic acid dye (pb11141, Beijing, China), WB APS (10002618, Shanghai, China); SDS (China Dalian Meilun Article No.: mb2479); TEMED (Aladdin, Shanghai, China, Article No.: t105497); Tween-20 (Sinopharm Shanghai, China, Article No.: 30189328); Acrylamide (Jintai Hongda, Beijing, China, art. No. 0341); Glycine (sigma v900144, USA); Methylene bisacrylamide (sigma, USA, art. No. 0172); Li Chunhong (China Shanghai Guoyao article number: 0172); Skimmed milk powder (pllay, Beijing, China, article number: p1622); BSA (Saibao, Yancheng, China); Ripa cracking liquid (biyuntian, Shanghai, China, Article No.: p0013b); Protease inhibitor (Jintai Hongda, Beijing, China, art. No.: 583794); Protein phosphatase inhibitor (pllai, Beijing, China, Article No.: p1260); Superecl plus (advansta, USA, Article No.: k-12045-d50); Developer (Shanghai Jiaxin Article No.: bw-61); Fixer (China Jiaxin Article No.: bw-62); Caspase-11 (Santa Cruz, USA, Article No.: sc-56038); Gsdmd-n (Abcam, UK, Article No.: ab219800); IL-6 (Abcam, UK, Article No.: ab233706); IL-1 β ( British Abcam, article number: ab205924);Mouse monoclonal antibody β- Actin (U.S. protein No.: 60008-1-ig); Goat anti rabbit second antibody (Article No.: sa00001-1); Goat anti mouse second antibody (sa00001-2); Rat TNF - α expression by ELISA- α ELISA reagent (Article No.: 88-7340)

### Apparatus

Fluorescent quantitative RCP instrument (thermo Article No.: pikoreal96, USA); Electrophoresis instrument (Beijing, China, Article No.: dyy-2c); Rotary film instrument (Beijing, China, Article No.: dycz-40d); Chemiluminescence imaging system A(Guangzhou Qinxiang article number: chemiscope 6100); Table type freezing centrifuge (Hunan Xiangyi, China, Article No.: h1650r); Automatic enzyme label washing machine (Shenzhen huisong Technology Development Co., Ltd. Article No.: pw-812); Multifunctional enzyme label analyzer (Shenzhen huisong Technology Development Co., Ltd. Article No.: mb-530)

## Methods

### Grouping and preparation of animal models

A rat model of COPD associated pulmonary hypertension was established by cigarette smoke exposure and lipopolysaccharide infusion (9). Sixty rats were randomly divided into normal group, western medicine group, model group, high-dose group, middle-dose group, and low-dose group, with 10 rats in each group. On the 1st and 14th day of the experiment, after pentobarbital sodium anesthesia, the model group and each drug group were instilled with LPS (1 mg/kg) intratracheally, and the normal group was not treated. No smoking was given on the same day. Except for the first day and the 14th day, the rats in each administration group and model group were placed in the self-made animal plexiglass fumigation box. Close the lid and seal the gap with plastic wrap to prevent fumes from escaping. Huangguoshu brand cigarette (10 mg flue-cured tar; Guizhou Zhongyan Industry Co., Ltd.) was inserted into the self-made tray, ignited it and put it into the fumigation box. 20 cigarettes were lit each time, once in the morning and afternoon. 2 hours each time (10 minutes apart). 6 days a week, a total of 60 days.

## Test index

### Detection of the expression of caspase-11, GSDMD and IL-6 in lung tissue by real time PCR

Take 0.25mg lung tissue, add 1ml Trizol, grind thoroughly, mix well, and split in the chamber for 5min. After adding 200ul of chloroform, shake vigorously for 15s, and let stand for 3min at room temperature. After centrifugation at 12,000 rpm for 10 min at 4°C, the supernatant was removed. And 1 ml of 75% sterile DEPC-treated water was added to wash the precipitate. Centrifuge at 12,000 rpm and 4°C for 3 min, discard the supernatant. Then air-dry for 5 min and add 20 UL of sterile non-enzymatic water to dissolve the precipitate. The RNA concentration (ng / UL) = a260 * Dilution Times * 40, RNA purity = od260 / od280. The acquired RNA sample was reverse tr.anscribed to acquire the corresponding cDNA. RNA samples were added into the nuclease free centrifuge tube and reverse transcribed to obtain the corresponding cDNA. The RNA sample was added into the nuclease free centrifuge tube, and then oligo (DT) 1 was added μL and random 1 μL DdH2O supplement. After heating (70 °C, 5 min), put it on ice to cool for 2 minutes. The reaction solution was gathered through centrifugation, and 4 UL of dtnp (2.5 mm each) was added × RT buffer4ul and primer mix; Add 200u / UL hifiscript and mix well. Two warm baths (25 °C for 10min; 2; The reaction was terminated by heating at 42 °C for 50 min and 80 °C for 10 min. 2 - △CT method was used for fluorescence quantitative analysis. The reaction conditions included 95 °C for 10 min, 95 °C for 15 s, 60 °C for 30 s and 40 cycles. The relative expression of the target gene was calculated by 2-ΔCT. The primers were synthesized by Shanghai Shenggong synthetic primers. The primer design is shown in Table 1.

**Table 1.**
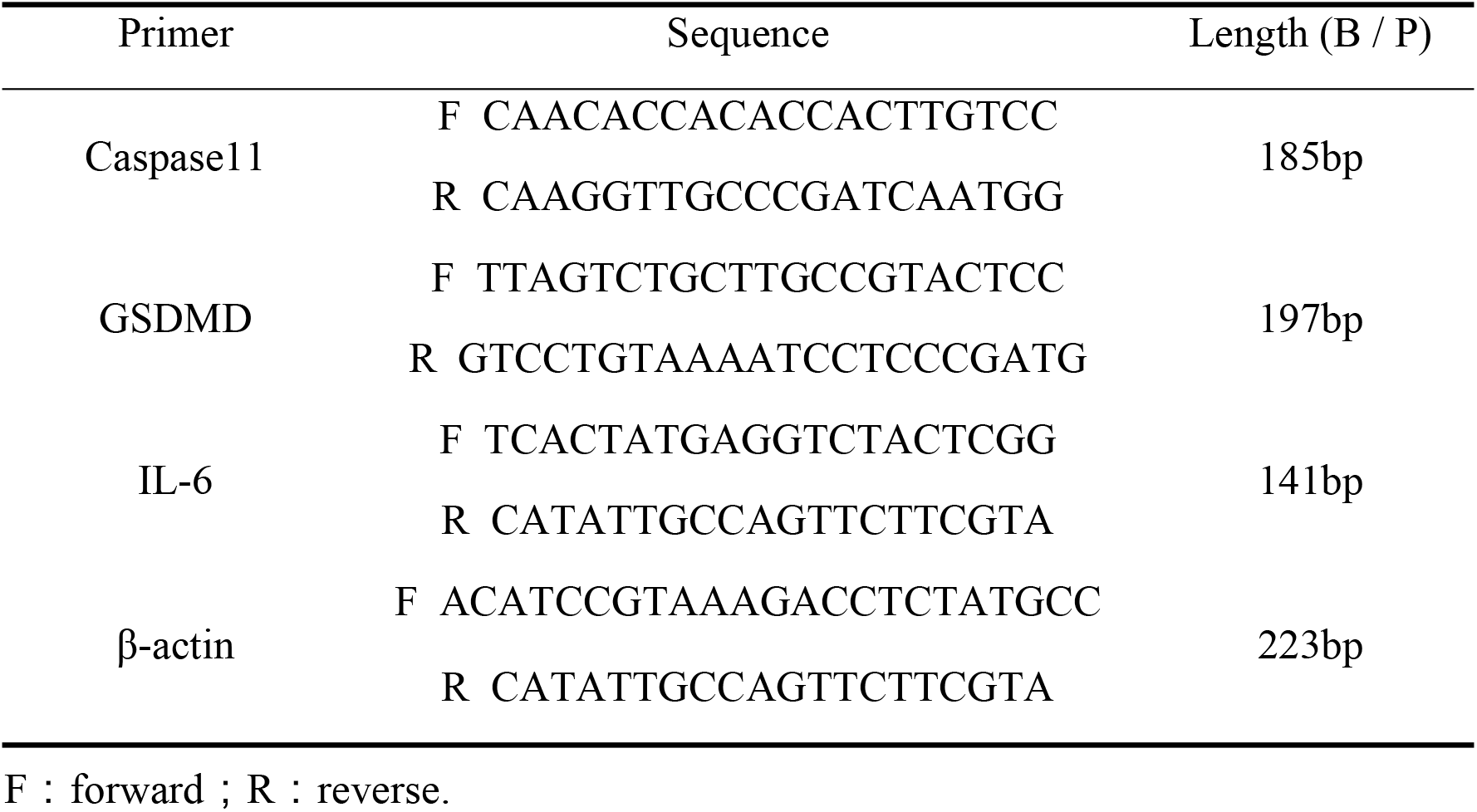
Primer sequence of PCR

### The expressions of Caspase-11, GSDMD and IL-6 proteins were detected by Western blot

After 2.5mg lung tissue of rats in each group was washed with ice pre cooled PBS. And put it into EP tube, then ripa lysate was added to grind it thoroughly. Then it was cracked on ice for 10min, centrifuged at 4 °C and 12000rpm for 15min, and take the supernatant. We used the BCA method to determine the protein concentration. After the protein concentration was determined, the protein was boiled for 12 min to denaturate. SDS-PAGE gel electrophoresis was prepared. With 20 μg protein was added to 10% polyamine gel, the protein mixture was separated by SDS-PAGE. Then transfer to PVDF model at 300mA for 130min, prepare 5% skim milk with PBST, seal for 90min, and wash PBST three times for 5min each time. Diluted I antibody was added, incubate the membrane with I antibody, and shake the bed overnight at 4 °C; TBST was washed 4 times for 10 min each, then horseradish peroxidase-conjugated II antibody was added and incubated for 90 min at room temperature. After incubation, TBST was washed 3 times for 10 min each. GAPDH was used as internal reference, ECL was used to develop and take pictures in the imaging system. The results were analyzed by quantity one software.

### Detection of TNF-α in serum through ELISA Level detection

A certain amount of lung tissue was collected and ELISA kit was used to determine TNF-α.The content of the product was determined.

### The morphology of lung tissue was observed by hematoxylin and eosin (HE) staining and the degree of pulmonary vascular muscle was determined by immunohistochemistry

After the experiment, the rat was killed, and the extracted lung tissue was fixed with 10% formaldehyde. After 1 week, the paraffin-embedded lung tissue was cut into 4 μm thick, baked at 60 °C and perform hematoxylin eosin (HE) staining. Lung tissue from each rat was prepared and placed under the microscope. Five high-power microscope fields were randomly selected for observation. Nikon ti-s inverted microscopes were used to analyze and process images. The expression of α-SMA in pulmonary blood vessels was detected by immunohistochemistry to evaluate the degree of pulmonary vascularization.

### Statistical analysis

The data were analyzed and processed by IBMS PSS Statistical 20.0 statistical software. The mean value was used in the experimental data ± Standard deviation 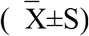. The LSD test was used when the variance was normally distributed, and Dunnett’s T3 test was used when the variance was not uniform. ANOVA was used to compare the data. The skew distribution was tested by rank sum test. Test level was set at both sides α =0. 05, P < 0.05 was considered statistically significant.

## Results

### The effect of compound tanglizi Decoction on caspase-11, GSDMD, IL-6mRNA expression in lung tissue of COPD PH rats

Compared with the normal group, caspase11, the expression of caspase11, GSDMD and IL-6 mRNA in the lung tissue of the model group was significantly increased (P<0.01); Compared with model group, the expression of caspase-11, GSDMD and IL-6mRNA in lung tissue of rats in the high-dose group, middle-dose group, low-dose group and western medicine group were significantly decreased. The results were shown in Table 2.

**Table 2.**
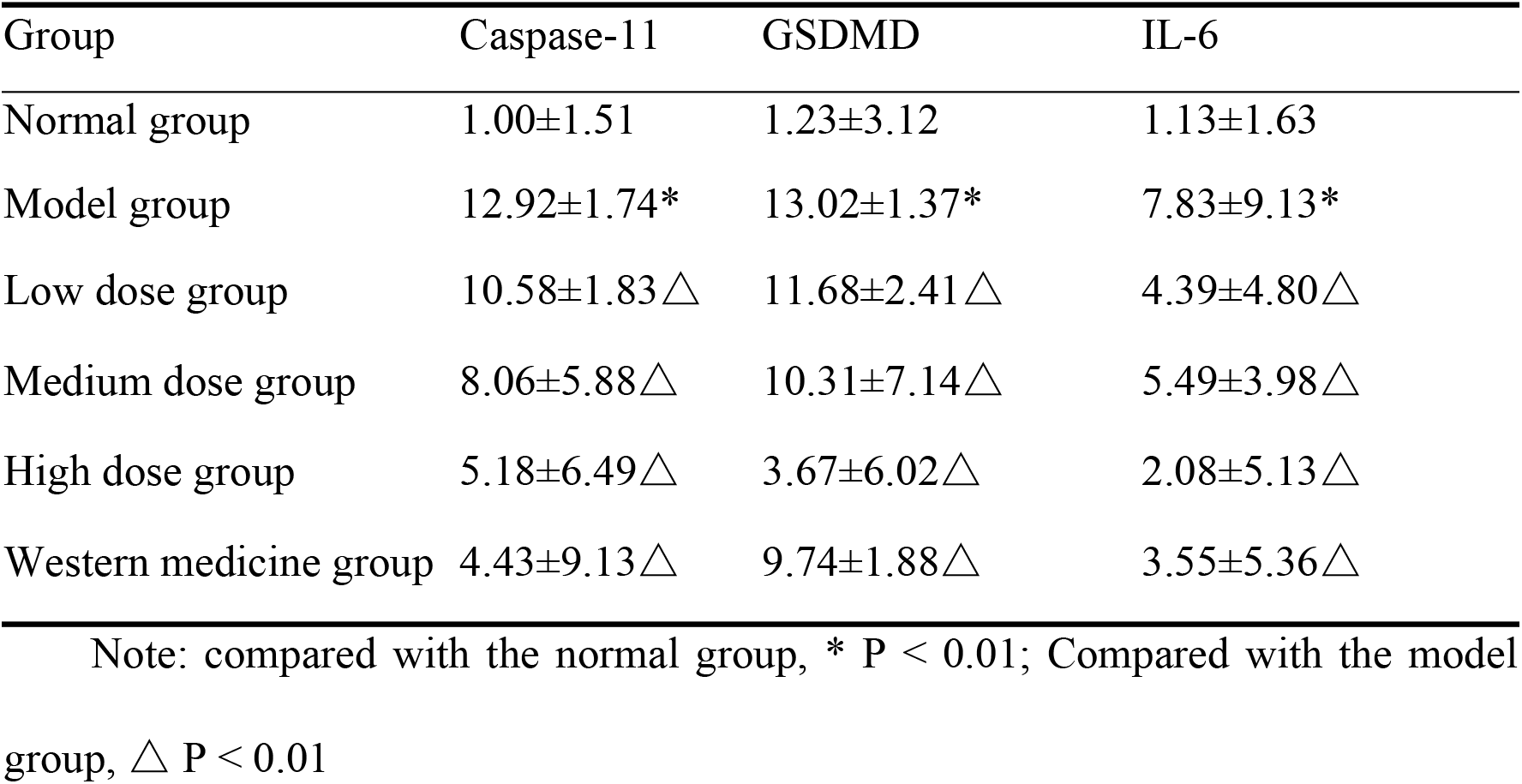
Effect of compound Tinglizi Decoction on expression of caspase-11, GSDMD and IL-6 mRNA in lung tissue of COPH-PAH rats(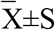, n=10)

### Effect of compound Tinglizi Decoction on the expression of caspase-11, GSDMD and IL-6 in lung tissue of COPH-PAH rats

WB results showed that compared with the normal group, the expressions of caspase-11(Figure 1 B), GSDMD(Figure 1 C) and IL-6(Figure 1 D) in the lung tissue of the model group were significantly increased (P < 0.01); Compared with the model group, the expressions of caspase-11(Figure 1 B), GSDMD(Figure 1 C) and IL-6(Figure 1 D) in high-dose, medium dose, low-dose and Western medicine groups of compound Tinglizi Decoction were significantly decreased (P < 0.01). The results were shown in Figure 1.

**Figure 1.**
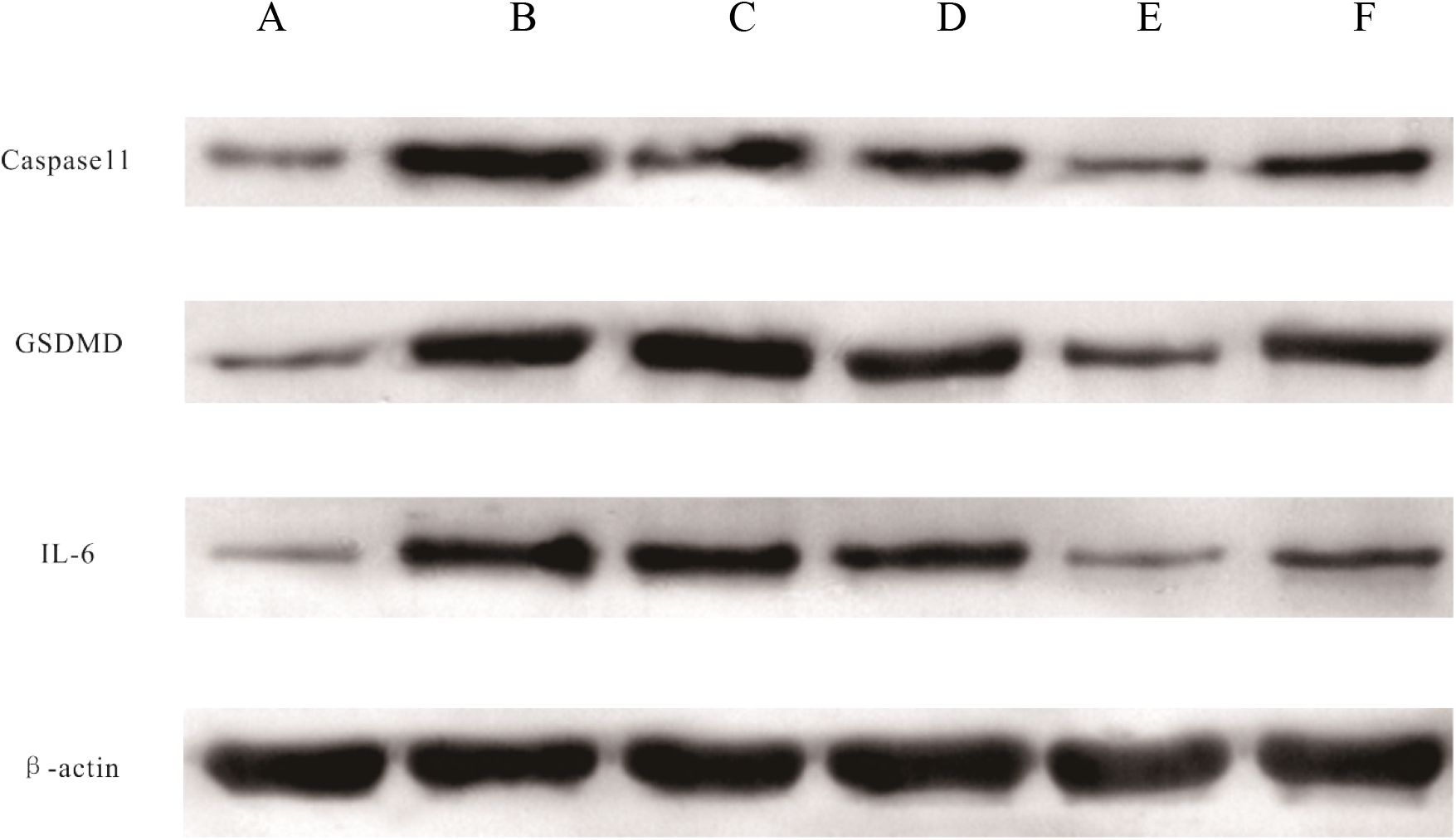
Western blot analysis.(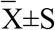, n=10)

### Compound decoction of triplizi in serum TNF- α of COPD PAH rats Level impact

The results showed that compared with the normal group, the TNF- α in the model group was significantly higher (P <0.01); the level of serum creatinine was significantly increased (P < 0.1); Compared with the model group, the levels of the low dose group, the middle dose group, the high dose group and Wetern group were significantly decreased (P < 0.05), and the results were shown in Table 3 and Figure2.

**Table 3.** Effect of compound Tinglizi Decoction on TNF-α levels in COPD-PAH rat serum (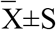, n=10)

| Group | TNF-α |
| --- | --- |
| Normal group | 33.78±9.85 |
| Model group | 30.56±7.85* |
| Low dose group | 38.17±10.85*△ |
| Medium dose group | 31.53±10.39*△ |
| Medium dose group | 34.27±8.99*△ |
| Western medicine group | 32.94±10.84*△ |

Note: Compared with the model group, △ $P < 0.01$ ; compared with the normal group, \* $P < 0.01$

**Figure 2.**
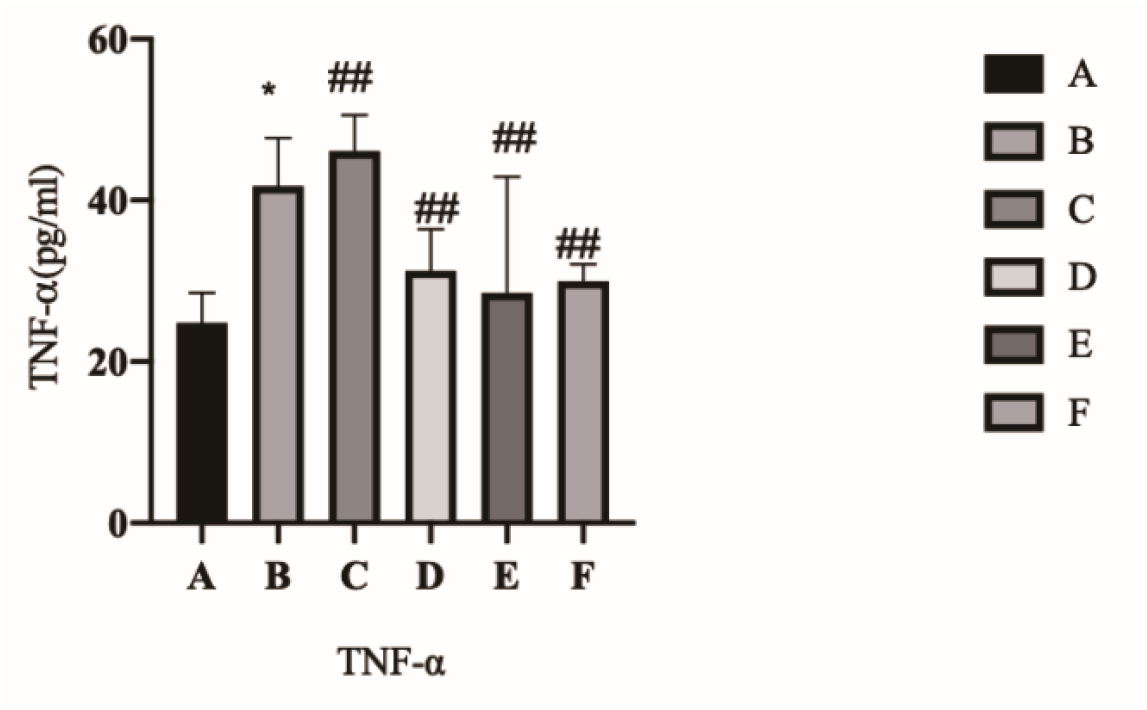
Effect of compound Tinglizi Decoction on TNF-α(a) levels in COPD-PAH rat serum (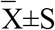, n=10)

### Pathomorphological observation of rats in each group

HE staining was performed in the lung tissue of rats. Under the microscope, the trachea and alveoli in the normal group were intact, and there was no obvious inflammatory cell exudation under the mucosa. After modeling, compared with the normal lung tissue, the model group showed damaged tracheal structure, exudation of a large number of inflammatory cells under the mucosa, atrophy of alveolar structure, and increase of microvessels. After drug intervention, compared with the model group, the western medicine group, the middle-dose group and the low-dose group showed a significant increase in the area of the organ cavity, good cell continuity, decreased tube wall thickening, and decreased inflammatory cells around the blood vessels. As is shown in Figure 3.

**Figure 3.**
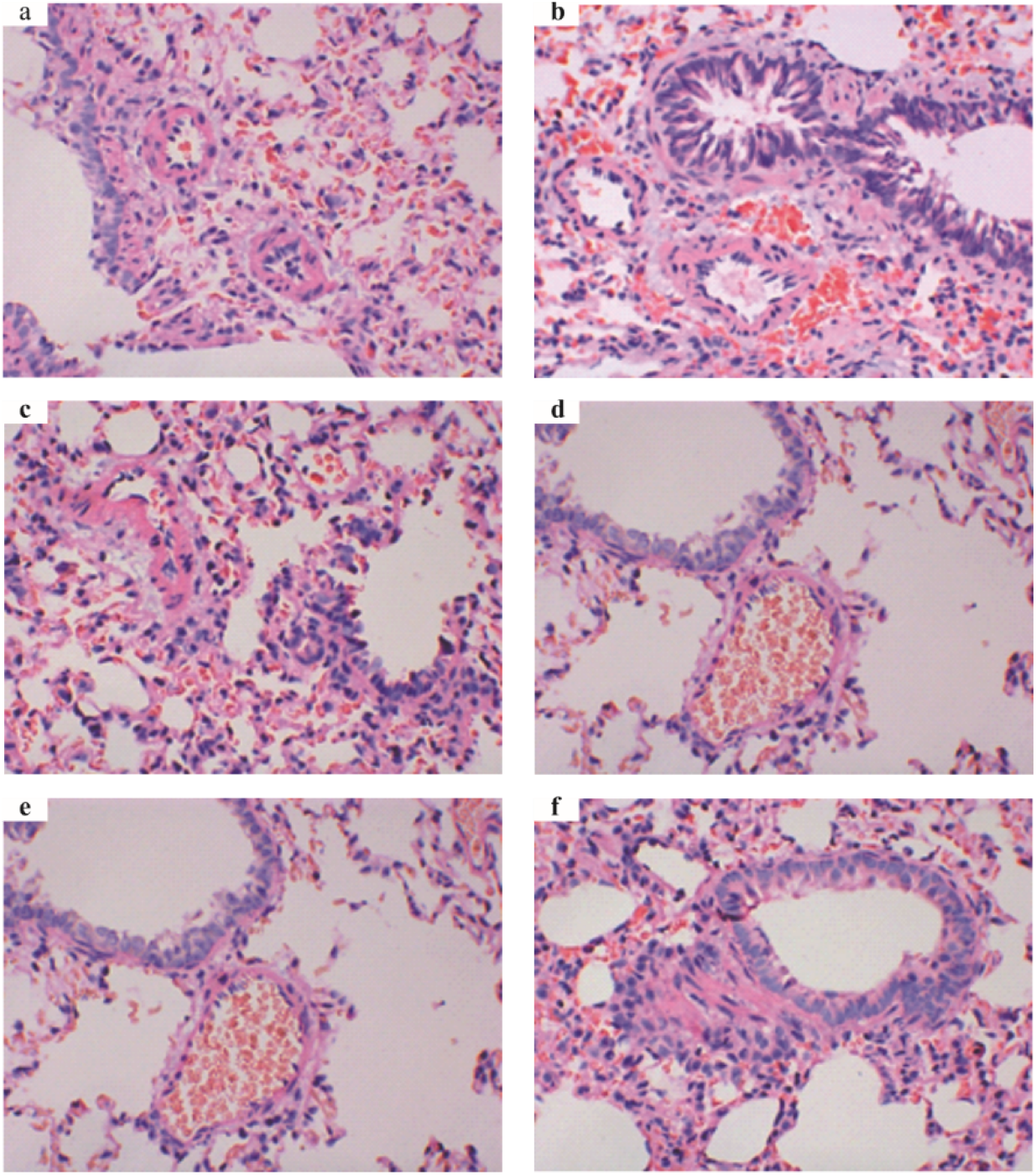
Pathomorphological observation of rats in each group, a: Normal group; b: Model group; c: Low dose group; d: Medium dose group; e: High dose group; f: Western medicine group.

**Figure 4.**
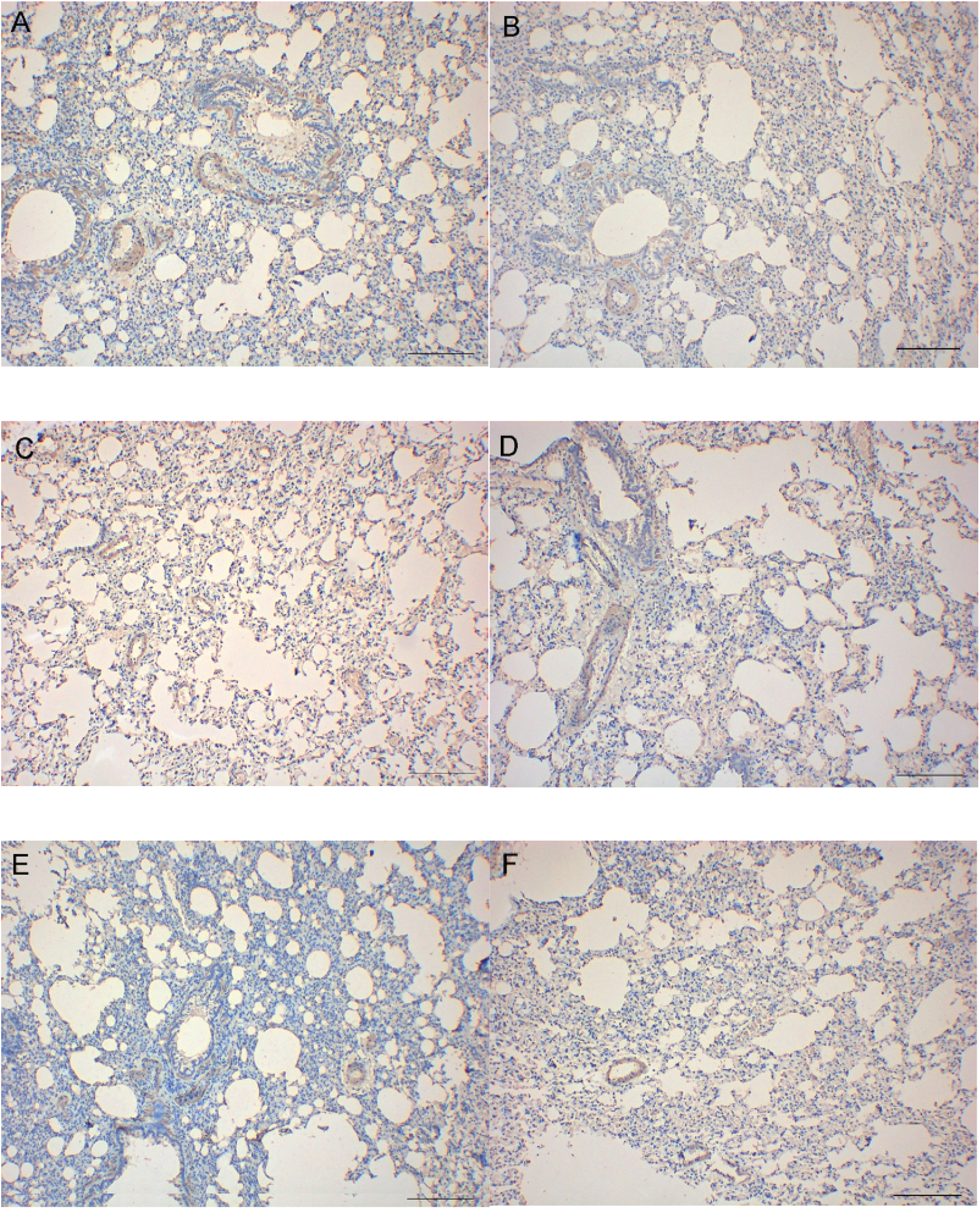
Pulmonary vascular muscle level (immunohistochemistry×100)

### Degree of vascularization

Immunohistochemistry showed that the rats in the blank control group were mainly non myogenic and partially myogenic vessels, and the fully myogenic vessels in the model group increased significantly. Compared with the model group, the vascularization of low-dose group, medium dose group, high-dose group and Western medicine group decreased in varying degrees.

## Discussion

The pathogenesis of pulmonary hypertension is complex. It is believed that it is related to inflammatory cell infiltration around the vascular lumen. The inflammatory lesions and the thickness of the vessel wall of the lesions are related to the mean pulmonary arterial pressure. This also makes us clear about the important role of inflammatory response in the process of pulmonary vascular remodeling (10). Pulmonary vascular remodeling is an important sign of COPD progression to pulmonary hypertension (11). The existing scholars believe that the pathogenesis of pulmonary hypertension is closely related to inflammation. Chemokines and cytokines can induce vascular remodeling to lead to PAH. Inflammatory cells and pulmonary vascular cells are the main sources of these two factors. Overexpression of these two factors can lead to the proliferation of vascular cells and increase their contractility. The abnormal increase of cytokines and chemokines in circulation leads to infiltration of immune cells around the lumen, including TNF- α, IL-1, IL-6, etc. IL-6 and other cytokines can regulate the abnormal proliferation, differentiation and migration of pulmonary vascular cells. The involvement of inflammatory cells is one of the main pathological mechanisms of PAH. Inflammatory response plays a key role in the pathogenesis of COPD associated pulmonary hypertension.

Pyroptosis is a programmed necrosis of cells mediated by GSDMD. Inflammatory bodies activate caspase, cut GSDMD to separate C-terminal and the N-terminal, breaking self-inhibition. Through the interaction of positive and negative charges, honeycomb-like pores are formed in the cell membrane, resulting in water conduction. Then the ion gradient inside and outside the cell membrane disappears, the cell swells and penetrates and dissolves, and finally the cell breaks down and dies. Many factors can cause cell focal death, including invasion of various pathogens, and the release of danger signals from external stimuli. Usually, focal death is a kind of favorable adaptive response initiated by host to deal with exogenous pathogens or endogenous risk signals. It can clear pathogens and damaged cells in vivo, and induce inflammatory response to resist infection. Focal death mediates the inflammatory response of COPD-PAH, thus inhibiting the occurrence of excessive focal death and becoming a new target for the treatment of pulmonary hypertension. The non-classical pathway is a focal death pathway which depends on caspase-4, 5 (human homology), and 11 (rat). Under the stimulation of bacteria and other signals, caspase-4, 5 and 11 are activated. Activated caspase-4, 5 and 11 can directly cleave GSDMD to form peptides containing N-terminal active domain, which can cause membrane perforation, increased permeability, release of cell contents, and inflammatory response. The occurrence of focal death of cells can lead to the release of inflammatory factors such as IL-6 and TNF-α, which can aggravate the inflammatory response of COPD-related pulmonary hypertension.

After long-term clinical practice, our team concluded that the deficiency of lung, spleen and kidney are the causes of pulmonary hypertension, and take the stagnation of phlegm and blood stasis as the standard. Treatment should be focus on nourishing the lungs, spleen, kidneys, removing blood stasis and removing phlegm (12). Compound Tinglizi decoction is a traditional Chinese medicine compound preparation founded by Professor Zheng-ping Bai under the guidance of the principle of “treating both yin and Yu”. It is composed of Tingli Dazao xiefei decoction and Linggui Zhugan Decoction. The composition of the prescription is: lepidium seed 15g, peach kerne l10g, carthamus tinctorious10g, leech 6g, ligusticum wallichii15g, poria cocos15g, cassia twig 15g, bighead atractylodes rhizome 15g, ardisia 10g, scutellaria baicalensis 10g, liquorice 6g and licorice 6g. The whole prescription is compatible with dispelling lung, resolving phlegm, relieving asthma. And in clinical verification (13), Compound TingliZi Decoction has the effects of reducing pulmonary arteriovenous pressure, improving blood circulation and reducing pulmonary congestion. However, the mechanism of how compound Tinglizi decoction can reduce inflammatory reaction and inhibit pulmonary vascular remodeling by inhibiting cell death is still unclear.

In the experiment, we found that the activation of non-classical pyrolytic pathway, the key proteins of pyrolytic, GSDMD, IL-6 and TNF-α, existed in COPD related pulmonary hypertension rats induced by cigarette smoke exposure combined with lipopolysaccharide. Compound Tinglizi decoction can inhibit the activation of non-classical cell death pathway and reduce the level of proinflammatory factors in COPD related pulmonary hypertension rats. This suggests that compound Tinglizi decoction may intervene COPD related pulmonary hypertension through non-classical way of regulating cell death.

## Conclusions

The mechanism of compound Tinglizi Decoction in the treatment of COPD-related pulmonary hypertension may be associated with the inhibition of non-classical pathway of cell death and the reduction of the level of pro-inflammatory factors, which provides a certain theoretical basis for the clinical treatment of COPD-related pulmonary hypertension.

## Acknowledgments

Not applicable.

## Availability of data and materials

The data used to support the findings of this study are available from the corresponding author upon request.

## Authors’ Contributions

Qi-zhi Wang and Yu Liu designed the study and were responsible for data acquisition. All authors contributed to the analysis and interpretation of data, writing, critical review, and approval of the final version.

## Ethics approval and consent to participate

All animal experiments followed stated protocols from the The Affiliated Hospital of Hunan Academy of Chinese Medicine and Hunan Shrek Jingda [permit to syxk (Xiang) 2019-0017.]

